# A single-cell RNA-seq catalog of ground truth gene coregulation

**DOI:** 10.64898/2026.09.18.752692

**Authors:** Valentine Svensson

## Abstract

Single-cell RNA-sequencing measurements are uniquely well-poised to identify coregulated gene transcription. Coregulation should be apparent as correlation of expression levels, but at which unit to quantify expression for calculation of correlations is not clear. To enable evaluation of normalization methods for identification of coregulation we have curated and preprocessed 35 scRNA-seq samples with known regulature relationships into a catalog. These datasets contain promoter-reporter data representing known positive coregulation and B cells where isotypic mutual exclusion of *κ* and *λ* light chains represent known negative coregulation. As an example, we use the catalog to evaluate seven normalization methods. Different normalization methods perform better for positive or negative coregulation, and results indicate implementation choices for advanced normalization methods The catalog may serve as a resource for evaluating novel proposed normalization methods and is available at https://huggingface.co/datasets/valsv/scrna-coregulation-benchmark

---

One goal of single-cell RNA sequencing (scRNA-seq) measurements is to learn transcriptional coregulation of genes (Su et al., 2023). Two genes being coregulated means that producing more expression of one genes also means producing more (or less) expression of the other gene, resulting in observed coexpression.

scRNA-seq captures noisy counts dominated by technical or uninteresting factors (Hicks et al., 2018). Normalization methods aim to represent biologically meaningful variation in gene expression, while accounting for technical factors. Strategies for normalizing data range from simple transformations to machine learning models that can output normalized units which quantitively reflects interpretable expression levels (Lopez et al., 2018; Lun et al., 2016).

Normalization methods are typically evaluated based on clustering or differential expression performance (Ge et al., 2025), but rarely on the ability to identify coregulation of genes across cells (Chau et al., 2025).

A major challenge with this type of data is the lack of ground truth. In the case of coregulation, we can focus on general statements considered true about transcriptional coregulation:

- In promoter-reporter systems, it is assumed that the fluorescent reporter expression correlates with endogenous gene expression (Chau et al., 2025).
- In B cells, kappa light chain genes are not expressed at the same time as lambda light chain genes due to isotypic exclusion (Vettermann & Schlissel, 2010) (except in rare cases (Giachino et al., 1995)).
- These correlations are assumed to be higher (or lower) than average correlation between two arbitrary genes. We can assume that, on average, correlation between these targets and arbitrary genes should have correlation 0 (in the absence of global uninteresting factors).

Normalization methods should reflect these biological truths. Here we present a catalog of preprocessed scRNA-seq data from these truth categories from 35 independent samples across 11 studies (Table 1). These can be used to evaluate normalization methods in their ability to reflect coregulation. We define how to measure consistency between biological truth and normalisation units using the catalog, and apply it to compare implementation options in SCVI normalization.

**Table 1:** Catalog overview. Studies are GEO series; samples are catalog files available as individual H5AD files.

| Species | Studies | Samples | Cells |
| --- | --- | --- | --- |
| **Promoter-reporter** |  |  |  |
| *Mus musculus* | 7 | 17 | 159,780 |
| *Arabidopsis thaliana* | 1 | 3 | 32,078 |
| **Isotypic exclusion** |  |  |  |
| *Homo sapiens* | 3 | 15 | 198,909 |
| **Total** | **11** | **35** | **390,767** |

The catalog is available at https://huggingface.co/datasets/valsv/scrna-coregulation-benchmark.

## Endognous gene and reporter transgene in promoter-reporter systems should manifest as positive correlation

Fluorescent reporter genes controlled by regulatory elements that have been introduced into cells enable quantification of transcriptional regulation using imaging and fluorometric technologies (Chalfie et al., 1994). In these systems, fluorescence signal from a reporter, such as eGFP or dsRed, are used as a proxy for gene expression.

In these promoter-reporter systems, expression of fluorescent reporters are driven by the promoter and additional regulatory elements which targets a corresponding endogenous gene. These systems are typically designed for fluorescence assays such as fluorescence microscopy or flow cytometry.

Cells engineered to express the reporters can also be measured through single-cell RNA-sequencing. If the promoter-reporter system maintains the corresponding endogenous gene, both transcripts from that gene, and from the gene encoding the reporter, can be quantified through molecule counts (Chau et al., 2025).

Since the promoter-reporter systems are designed to express the reporter in the same context as the endogenous gene is expressed, the relation between them should manifest as strong positive correlation.

### On average, other genes should have zero correlation with the fluorescent reporter gene

While some genes may be coexpressed with the fluorescent reporter, most genes are not. After normalizing out global, uninteresting, effects, a random sample of genes should on average have no correlation with the fluorescent reporter gene.

Optimally, a normalization of expression levels should be close to the (0, 1) when measuring correlation between the reporter and background genes on the x-axis versus the reporter and the corresponding endogenous gene on the y-axis.

## Mutual isotypic exclusion of the IGK locus or IGL locus in B cells should manifest as negative correlation

During development of B cells, pre-B cells with functional heavy chains undergo rearrangement of the *κ* light chain locus (Igk, consisting of 46 functional genes for humans). If Igk rearrangement fails to produce a functional light chain, the Igk locus is silenced. Instead, the *λ* light chain locus (Igl, consisting of 42 genes for human) is opened and rearranged (Vettermann & Schlissel, 2010).

Pre-B cells without either functional *κ* or *λ* light chains do not develop into mature B cells and do not exit the bone marrow. B cells outside the bone marrow must express either Igk or Igl genes, but not genes from both loci (except in rare cases (Giachino et al., 1995))

Because the light chain loci are recombined during maturation, short read alignment to a reference is challenging. However, constant region genes stay true to the reference (IGKC for Igk, IGLC1-3 for Igl), allowing mapping and UMI counting in single B cells.

The mutual exclusivity of IGKC and IGLC genes means that the relation between these genes should manifest as negative correlation in expression level quantification.

### On average, other genes should have zero correlation with IGKC or IGLC1-3 in B cells

The IGKC and IGLC1-3 genes are part of the B cell program consisting of many coregulated genes. Individual cell will use this program to different extents. However, most genes will not be part of this program, and a randomly selected gene is not expected to correlate with the B cell program. A gene expression quantification should reflect an average correlation of zero with arbitrary background genes.

## Comparizon of normalization methods

We use the catalog and the two coregulation truth classes to evaluate seven normalization methods. For each sample, normalization was performed by each of the seven normalization methods (Raw counts, log10(CPM + 1), log10(CPK + 1), PFlogPF, and three variants of SCVI normalization).

We calculated correlations using the normalized values for each sample, resulting in a pair of target correlation and average background correlation values for each sample, normalization - pair. To summarize results with a single value per normalization method, we calculated the mean distance to an optimal point (1.0 target correlation, 0.0 average background correlation for promoter-reporter coregulation, Figure 1 −1.0 target correlation, 0.0 average background correlation for isotypic exclusion, Figure 2).

**Figure 1:**
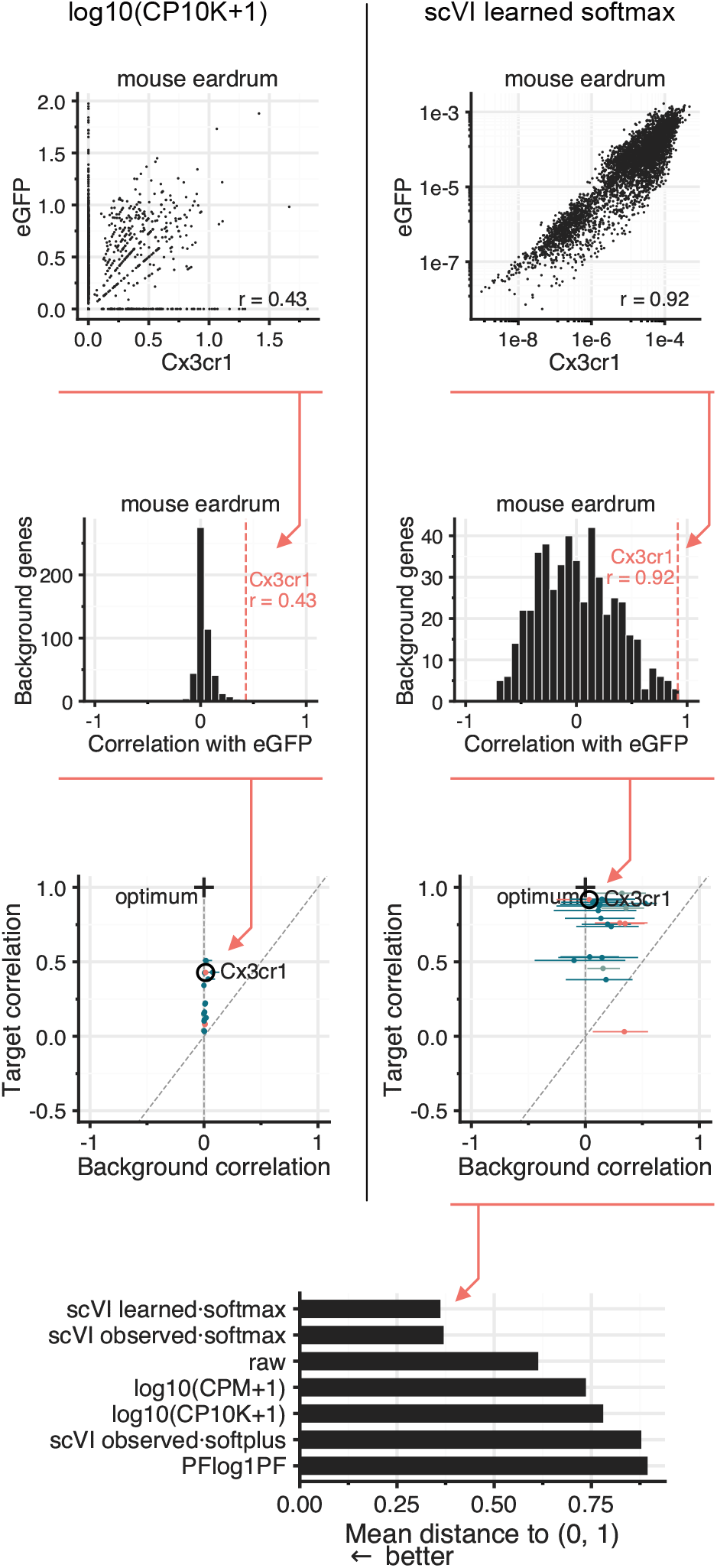
Evaluating normalization methods with promoter-reporter data. Correlations between the fluorescent reporter and the corresponding endogenous gene as well as correlation with background genes are calculated for each sample. Each normalization method is then evaluated as average distance to optimal performance.

**Figure 2:**
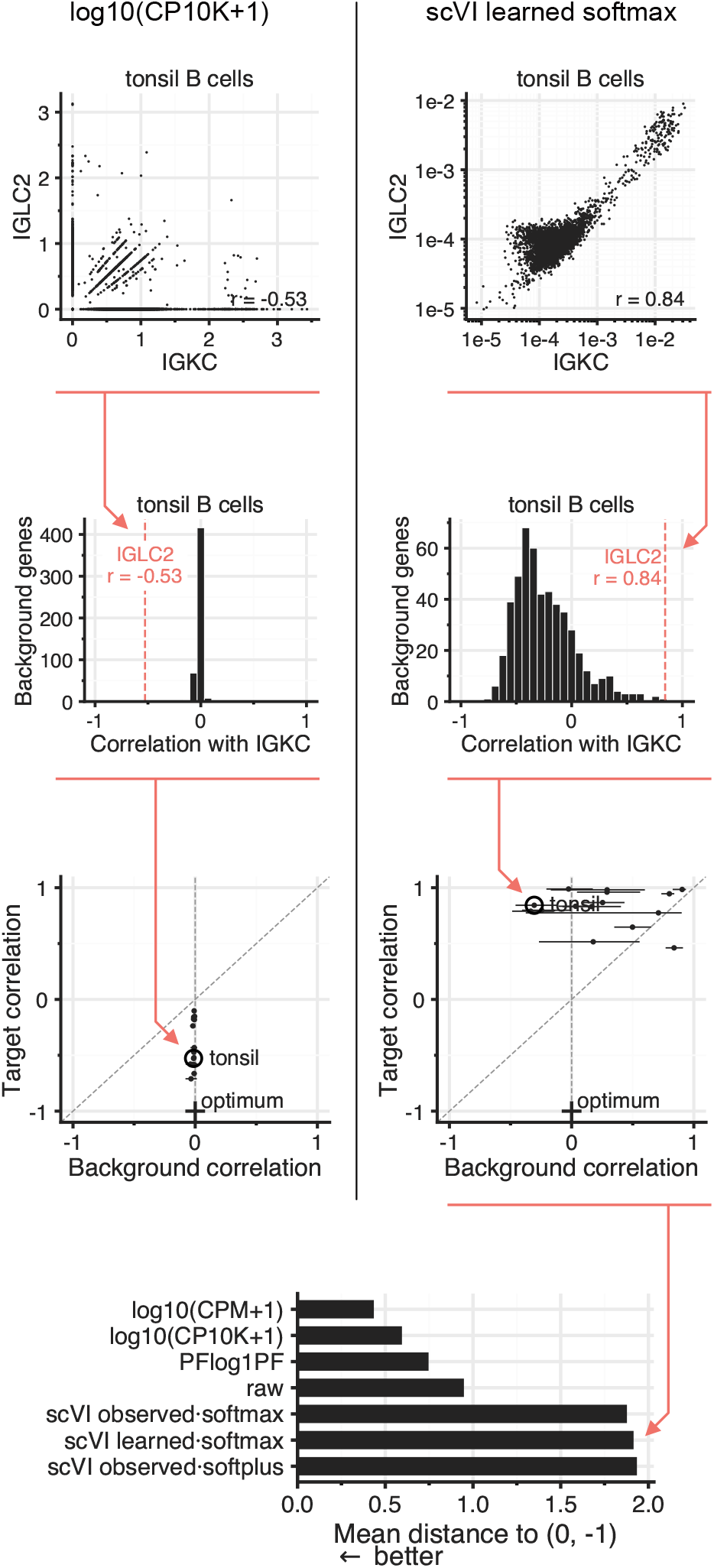
Evaluating normalization methods using isotypic exclusion data. Correlation between IGKC and IGLC2 or IGLC3 is calculated for B cells from each sample, as well as average correlation with background genes. The average distance to an ideal combination of mutual exclusive correlation and background correlation is used to quantify performance of a normalization method.

The normalization methods ranked differently in the positive coregulation and negative coregulation benchmarks.

The three SCVI variants used for normalization differed in whether they used observed library size or learned library as a latent variable, and whether gene expression decoding used a softmax ot softplus final activation function. The SCVI variants using softmax activation function had the highest performance at inferring positive coregulation (Figure 1). When using the softplus activation function, SCVI normalization induced spurios correlations with background genes.

Non-SCVI normalization methods showed near-zero background correlation, but less strong positive correlation between fluorescent reporter and corresponding endogenous gene. Un-normalized raw UMI counts performed better than standard log10(CPM + 1) or log10(CPK + 1) normalization.

In the negative coregulation benchmark based on isotypic exclusion, all three SCVI variants showed particularly poor performance, often inducing *positive* correlation between *λ* and *κ* genes (Figure 2).

For isotypic exclusion, the log10(CPM + 1) and log10(CPK + 1) normalizations performed better at identifying negative coregulation.

## Discussion

Unlike many scRNA-seq analysis tasks, these coregulation identification tasks have relations that are (mostly) considered to be truths which should be represented in data.

The assumed truths are not perfect or absolute. Different promoter-reporter systems will have different levels of correspondence between the fluorescent reporter and the endogenous gene it is reporting for. There are many parts of endogenous transcriptional regulation not reflected by an exogenous fluorescent reporter with a designed promoter sequence.

Similarly, regulation of expression of *κ* and *λ* light chain genes in B cells is well understood, but as with many aspects of biology, there are rare exceptions where mutual isotypic exclusion does not hold.

Using the evaluation metrics allows selecting between seemingly arbitrary implementation choices for model-based normalization to better reflect biological expectations of quantifications.

In some cases, performing no normalization at all might be the best choice for accurate inference.

These regulatory properties, largely considered to be true, should also be reflected when single-cell data is created by generative machine learning models. If used to generate B cell data, isotypic exclusion should be observable in the cells being generated.

By collecting this catalog, we hope it should be easier for researchers to test new proposed normalization methods. It can also be used to collect additional data on supposed true regulatory relationships.

## Methods

### Measuring gene coregulation

We evaluate normalized expression using gene pairs with an expected direction of coregulation. A fluorescent reporter and the endogenous gene associated with its regulatory elements should show positive correlation. In B cells, expression of the kappa and lambda immunoglobulin light chains should show negative correlation. These relationships provide reference directions without requiring the expression of either gene to be known in absolute units.

For each sample and normalization method, we calculate Pearson’s correlation across cells between the two genes. We use the reporter as the reference gene for promoter-reporter pairs, and *IGKC* as the reference for light-chain pairs. Reporter analyses include cells with zero reporter counts. For light-chain analyses, we select cells with at least one raw UMI assigned to *IGKC, IGLC2*, or *IGLC3*, and calculate correlations of *IGKC* with *IGLC2* and *IGLC3* separately. This selection is applied to both PBMC and sorted tonsil samples, so the evaluated correlations are conditional on detectable light-chain expression.

### Background correlations

A positive correlation between a reporter and its target can also arise from variation that affects many genes (Su et al., 2023). We therefore compare each target correlation with correlations between the same reference gene and a random set of background genes. This measures whether the expected relationship is recovered in the context of a general shift in gene-gene correlations.

For each sample and pair, we sample up to 500 background genes without replacement using a NumPy random generator initialized with seed 42. Genes must have nonzero raw counts in at least 10 cells. For the catalog summaries, this threshold is evaluated within the population used to calculate correlations. We exclude all reporter features and the current endogenous target from promoter-reporter backgrounds, and genes with symbols beginning with IGK, IGL, IGH, or IGLL from light-chain backgrounds. The three scVI configurations share a background set, but background sets are not fixed across all normalization methods.

We summarize background correlations by their median and interquartile range, omitting undefined correlations from constant expression vectors. Individual background genes can be biologically coregulated with the reference; the distribution provides an empirical baseline for interpreting the target pair. Its interquartile range describes variation among background genes, rather than uncertainty across cells or biological replicates.

### Distance from the reference relationship

We represent each sample-gene pair by its median background correlation *b* and target correlation *r*. A normalization that preserves the expected relationship while centering background correlations near zero approaches (0, 1) for promoter-reporter pairs or (0, −1) for light-chain pairs. We define the distance from this reference point as

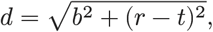

where *t* = 1 for promoter-reporter pairs and *t* = −1 for light-chain pairs. A smaller distance indicates a closer match to both criteria. The reference points express the desired correlation directions; they do not assume that every biological pair can attain perfect correlation.

We average distances with equal weight per sample-gene pair. The promoter-reporter summary contains 21 pairs, and the light-chain ranking contains the 15 *IGKC*-*IGLC2* pairs. *IGLC3* correlations are included in the accompanying tables. A study therefore contributes in proportion to its number of included pairs. The illustrative panels show eGFP and *Cx3cr1* in GSE296504 and *IGKC* and *IGLC2* in fraction A of GSE260943. These panels compare log10(CP10K + 1) with the scVI softmax configuration using an explicit observed size factor; the rankings include all seven normalization methods and configurations.

### Count normalization

Let *X*_*ig*_ be the UMI count of gene *g* in cell *i*, and *L*_*i*_ = ∑_*g*_ *X*_*ig*_ the total number of UMIs in that cell. We use raw counts as a baseline and compare two transformations that scale cells to a common total before taking the logarithm:

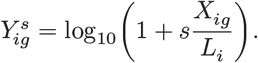

We set *s* = 10^4^ for log10(CP10K + 1) and *s* = 10^6^ for log10(CPM + 1). The pseudocount is added after scaling, so the choice of scale changes the relative contribution of low and high counts to the transformed values. Library totals include all genes in the input matrix.

We also evaluate PFlog1PF, which scales counts before and after a logarithmic transformation (Booeshaghi et al., 2022). First, we scale each cell to the mean library size 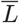 and calculate

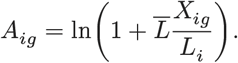

The logarithm changes each cell’s total. We calculate the new total *Q*_*i*_ = ∑_*g*_ *A*_*ig*_ and rescale the transformed values to its mean across cells, 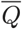:

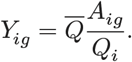

Both scaling steps use the complete gene matrix. We calculate Pearson correlations directly from the resulting values, without taking another logarithm. For the light-chain analysis, the PFlog1PF means and scaling factors are calculated within the selected population.

### scVI expression estimates

We use scVI with a negative-binomial observation model whose mean depends on a cell’s latent representation and an abundance scaling factor (Gayoso et al., 2022; Lopez et al., 2018). We fit a separate model to each sample and compare three configurations: a standard softmax decoder without an explicitly registered size-factor field, a softplus decoder with the observed UMI total registered as a size factor, and a softmax decoder with the same explicit size factor. Softmax constrains the decoder output to sum to one across genes, whereas softplus produces positive values without this constraint. The comparison therefore tests how decoder parameterization and the treatment of cell abundance affect the recovered correlations.

Each model uses 32 latent dimensions, a batch size of 512, a learning rate of 0.004, and a maximum of 200 training epochs. We set the maximum weight of the Kullback-Leibler regularization term to 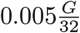, where *G* is the number of input genes. This scales its weight with the number of genes relative to the latent dimension. Models use the full input feature set, with no highly variable gene selection or additional batch or covariate fields. Reporter models are fit to their source count matrices, with the GSE160772 cell filter applied before training; human models are fit to the harmonized sample matrices described below.

We extract expression with `get_normalized_expression` and apply a log10 transformation before calculating correlations. Human models are trained on the full sample, and their expression estimates are subsequently restricted to cells passing the light-chain selection. Before calculating scVI correlations, we omit rows with nonfinite log-expression values among the requested genes.

### Data

The catalog contains 35 sample matrices from 11 GEO studies (Table 1). Each sample corresponds to a capture or sample-specific matrix and does not necessarily represent an independent donor. We evaluate expression relationships within samples, without testing differences between treatment or disease groups.

The promoter-reporter data comprise 17 mouse samples and three *Arabidopsis thaliana* root samples. Mouse studies pair eGFP with *Pdgfrb* in endometrium (GSE160772 and GSE198556) (Kirkwood et al., 2021; Kirkwood et al., 2022), *Rorc* in large-intestinal lamina propria (GSE181864) (Zhou et al., 2022), *Il23r* in small intestine (GSE229976) (Ahmed et al., 2024), *Cx3cr1* in tympanic membrane (GSE296504) (Zhang et al., 2026), *Dlx1* in embryonic medial ganglionic eminence (GSE316394) (Molero et al., 2026), and *Sox9* in liver (GSE319345) (Kanakanui et al., 2026). GSE296504 additionally pairs DsRed with *Cspg4*, giving 21 pairs across the 20 samples. The Arabidopsis samples pair GFP reporters with *WER, CORTEX*, and *SCR* (GSE295703) (Chau et al., 2025). Reporter-target assignments follow the experimental constructs, which include knock-ins, BAC transgenes, and promoter fusions.

The light-chain data comprise 15 human samples from three studies. GSE306378 contains PBMCs from three healthy donors and three donors with systemic lupus erythematosus (Cheng et al., 2026). GSE285843 contains PBMCs from three donors profiled with both 10x and Parse technologies (Espinoza et al., 2026). GSE260943 contains three sorted B-cell fractions from one human tonsil donor (McGrath et al., 2025). These samples contribute 30 pairs when *IGLC2* and *IGLC3* are evaluated separately.

### Reporter quantification and count processing

To measure reporter transcripts alongside endogenous genes, we reprocess the seven mouse studies with kallisto and bustools through `kb count` using reporter-augmented references (Melsted et al., 2021). This is necessary when the original reference omits the reporter, as for eGFP in GSE181864. The shared reference uses GRCm39 with Ensembl release 115 annotation and a 720-nucleotide eGFP sequence represented as a single-exon gene. GSE296504 uses a separate GRCm39 reference with Ensembl release 113, eGFP (GenBank U55762.1), and mammalian codon-optimized DsRed (MN623118.1). We use the codon-optimized DsRed sequence to recover reporter reads that did not map to the wild-type *Discosoma* sequence.

The Parse Evercode WT v1 data in GSE319345 are processed using an adaptation of the Mortazavi laboratory Parse workflow, including collapse of reverse-transcription barcode representations before conversion to sample matrices. Arabidopsis and human data are imported from existing study-derived count matrices.

We retain barcodes with more than 500 UMIs in GSE160772 and all four GSE319345 samples, and at least 100 UMIs in GSE198556, GSE181864, and GSE296504. The GSE316394 samples GSM9451641, GSM9451642, and GSM9451643 use a threshold of more than 500 UMIs. Reporter detection is not an additional retention criterion.

We harmonize gene identifiers and feature order within each catalog, retaining reporter features and their endogenous-target assignments. Standard mouse matrices contain 78,335 features. We match GSE296504 by Ensembl identifier after removing version suffixes, represent 95 absent reference features with zeros, and append DsRed. Arabidopsis matrices retain 32,834 features. Human matrices use the 33,694-feature reference from GSE306378; we match GSE285843 by symbol or Ensembl identifier and GSE260943 by Ensembl identifier. Unmatched source features are omitted, and absent reference features are represented by zeros: 7,703 in GSE285843 and 1,343 in GSE260943. We recalculate library totals after harmonization.

### Data availability

The count matrices, sample and gene metadata, and correlation summaries are available at https://huggingface.co/datasets/valsv/scrna-coregulation-benchmark. Matrices are provided in AnnData format (Virshup et al., 2021), with reporter-target and light-chain pairs organized into separate catalogs.

